# *In vitro* characterization of protein effector export in the bradyzoite stage of *Toxoplasma gondii*

**DOI:** 10.1101/2020.01.08.899773

**Authors:** Joshua Mayoral, Peter Shamamian, Louis M. Weiss

## Abstract

The ubiquitous parasite *Toxoplasma gondii* exhibits an impressive ability to maintain a chronic infection of its host for prolonged periods. Despite this, little is known regarding if and how *T. gondii* bradyzoites, a quasi-dormant life-stage residing within intracellular cysts, manipulate the host cell so as to maintain a persistent infection. A previous proteomic study of the cyst wall, an amorphous layer of proteins that forms underneath the cyst membrane, identified MYR1 as a putative cyst wall protein *in vitro*. As MYR1 is known to be involved in the translocation of parasite derived effector proteins into the host cell, we sought to determine whether parasites transitioning toward the bradyzoite life stage retain the capacity to translocate proteins via this pathway. By epitope tagging the endogenous loci of four known effectors that translocate from the parasitophorous vacuole into the host cell nucleus, we show by immunofluorescence that most effectors accumulate in the host nucleus at early but not late timepoints post-infection during the tachyzoite to bradyzoite transition and when parasites farther along the bradyzoite differentiation continuum invade a new host cell. We demonstrate that the suppression of interferon-gamma (IFN-γ) signaling, previously shown to be mediated by the effector TgIST, also occurs in the context of prolonged infection with bradyzoites, and that TgIST export is a process that occurs beyond the early stages of host cell infection. These findings have important implications as to how this highly successful parasite maintains a persistent infection of its host.

**IMPORTANCE:** *Toxoplasma* bradyzoites persist within tissue cysts and are refractory to current treatments, serving as a reservoir for acute complications in settings of compromised immunity. Much remains to be understood regarding how this life-stage successfully establishes and maintains a persistent infection. In this study, we investigated whether the export of parasite effector proteins into the host cell occurs during the development of *in vitro* tissue cysts. We quantified the presence of four previously described effectors in host cell nuclei at different timepoints post-bradyzoite differentiation and found that they accumulate largely during the early stages of infection. Despite a decline in nuclear accumulation, we found that one of these effectors still mediates its function after prolonged infection with bradyzoites and provide evidence that this effector is exported beyond early infection stages. These findings suggest that effector export from within developing tissue cysts provides one potential mechanism by which this parasite achieves chronic infection.

## INTRODUCTION

The intracellular parasite *Toxoplasma gondii* is estimated to infect up to one-third of the global human population (1). The success of this pathogen can be partially attributed to its flexible life cycle, one in which a wide variety of hosts can be infected and transmit the latent life-stage following predation by another organism (2). During acute infection, the tachyzoite life-stage replicates quickly and robustly, disseminating to various tissues of the body (3). Although the host immune response can typically clear the majority of tachyzoites and overcome acute infection, a subset of parasites differentiate into the slowly growing life-stage termed the bradyzoite, which persists into chronic or latent infection (4). Bradyzoites persist within their host for an indefinite period of time, residing predominately in muscle tissue and the brain (4). The mechanisms by which bradyzoites manage to evade the host immune system and manipulate their host cell to optimize survival remain unclear and understudied.

Within the host cell, bradyzoites reside in a specialized vacuole termed the tissue cyst. Tissue cysts are typified by an amorphous collection of proteins and glycoconjugates termed the cyst wall, which forms underneath the limiting membrane, or cyst membrane, of the tissue cyst (5). Proteomic studies of the cyst wall from *in vitro* induced tissue cysts identified Myc-Regulating Protein 1 (MYR1) as a putative cyst wall protein (6). MYR1 has been previously shown to be involved in the process of parasite protein translocation from the parasitophorous vacuole into the host cell during tachyzoite infection (7). Thus far, exported effector proteins that have been identified to be secreted in this manner include GRA16 (8), GRA18 (9), GRA24 (10), TgIST (11, 12), and HCE1/TEEGR (13, 14). In the context of tachyzoite infection, these effectors have been shown to bind to host cell proteins, translocate into the host nucleus (GRA18 remains in the host cytoplasm), and affect host cell signaling pathways to favor the parasite (15). GRA28 is an additional protein identified by proximity-based biotinylation that has been shown to localize to the host cell nucleus during tachyzoite growth conditions (16), although the host cell targets of this protein are yet to be identified.

Given the presence of MYR1 protein in the cyst wall, we hypothesized that bradyzoites retain the capacity to continually export proteins from within tissue cysts, so as to maintain constant control of their host cell. We set about testing this hypothesis by first confirming the cyst wall localization of MYR1. After GRA16, GRA24, GRA28, and TgIST were epitope tagged at their endogenous loci, we found by immunofluorescence that the intensity of these effectors in the host nucleus declines over time in human fibroblasts containing individual vacuoles with differentiating parasites. A similar pattern of export was observed in fibroblasts infected with parasites farther along the bradyzoite differentiation continuum, as well as during infection of mouse primary cortical neurons. We found that renewed nuclear effector accumulation did not occur during a tachyzoite superinfection in host cells containing older tachyzoite or bradyzoite vacuoles, indicating that declining levels of ROP17 are likely not the cause for the observed pattern of effector export. Despite the observed decline of effector export, an inducible knockout approach revealed that host nuclear TgIST-3xHA is not detectable during the later phases of infection when deleted 1 day post-infection indicating that TgIST export continues to occur beyond day 1 post-infection.

## RESULTS

To validate the localization of MYR1 at the cyst wall of *in vitro* tissue cysts, the endogenous locus of MYR1 was epitope tagged at the C-terminus with 3 copies of the hemagglutinin tag (3xHA) in the PruΔku80 strain using CRISPR/Cas9 (Fig. 1A). After obtaining clonal populations of endogenously tagged parasites, immunofluorescence assays (IFAs) were performed on infected human foreskin fibroblast (HFF) cultures. Egressed tachyzoites were used for initial experiments, with bradyzoite differentiation induced at the time of invasion by replacing growth media with low serum alkaline media prior to the addition of parasites, followed by subsequent culture in an ambient CO_2_ incubator. MYR1 was detected in the nascent cyst wall of vacuoles undergoing differentiation as early as 1 day post-infection (p.i.) and out to 7 days p.i., as determined by colocalization with glycosylated CST1 (Fig. 1B), an early marker of bradyzoite differentiation (17). MYR1 cyst wall localization was also evident at time points between 2 to 6 days p.i. (Fig. S1). To determine whether cysts formed by parasites farther along the bradyzoite differentiation path also contained MYR1, egressed parasites cultured *in vitro* under bradyzoite inducing conditions for 6 days were used to infect a new HFF monolayer. Using GFP as a marker of bradyzoite differentiation (by virtue of GFP expression driven by the bradyzoite specific LDH2 promoter in the PruΔku80 strain (18)), MYR1 was shown to be expressed and secreted by these *in vitro* derived bradyzoites upon infection of new host cells under bradyzoite inducing conditions, both at 1 and 4 days p.i. (Fig. 1C).

**Figure 1.**
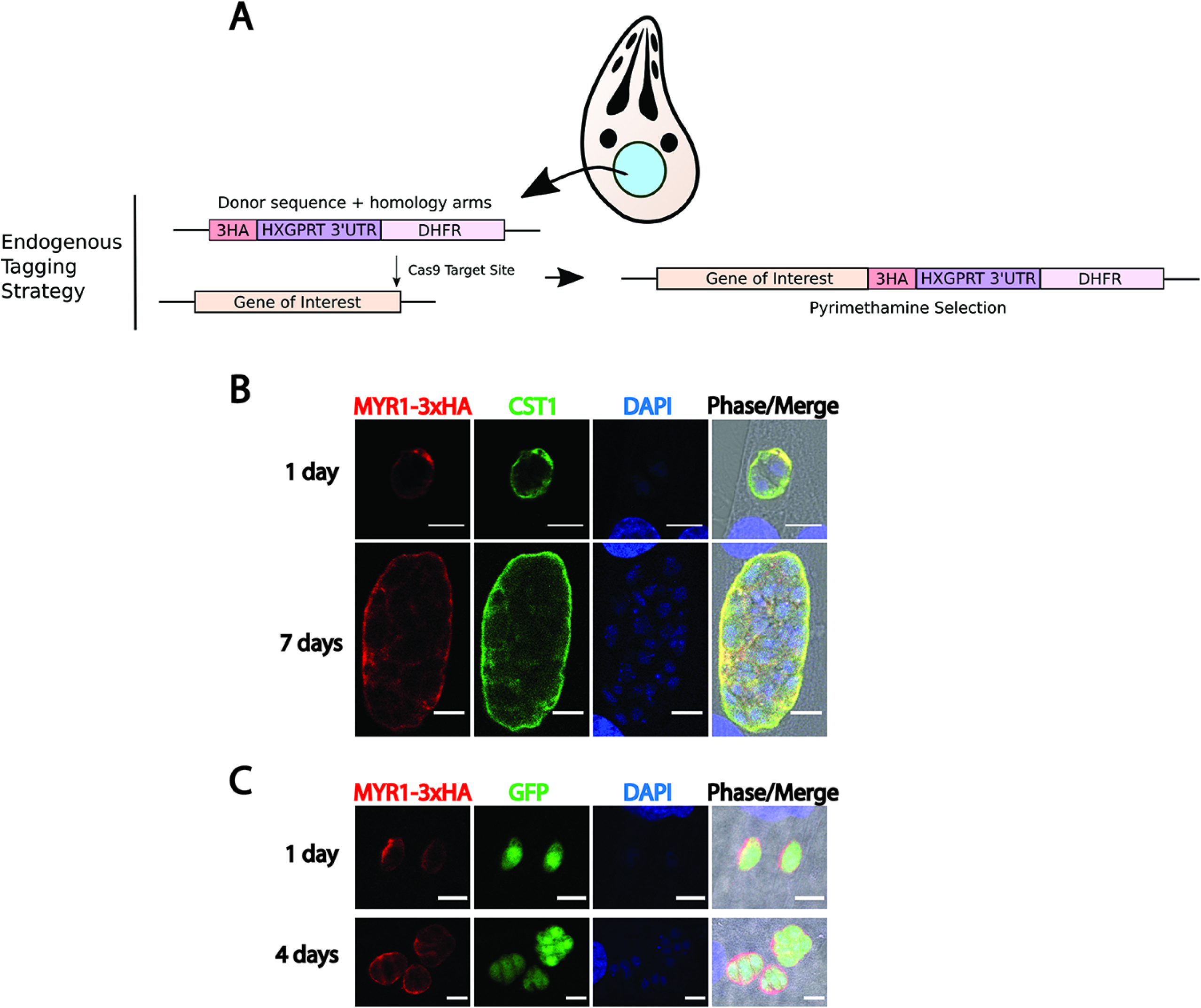
Endogenous tagging and immunofluorescence of MYR1-3xHA reveal MYR1 is secreted into the vacuole during bradyzoite differentiation and post-bradyzoite infection. (A) Diagram illustrating Cas9 strategy to introduce a 3xHA epitope tag, HXGPRT-3’UTR, and DHFR resistance cassette at the C-terminus of the gene of interest. Note that the Cas9 target site was located either in the C-terminus or 3’UTR of a gene, and the donor sequence containing at least 30bp of homology to endogenous sequences on both ends was amplified by PCR. See Supplementary Table S1 for a list of primers used. (B) Representative images of MYR1-3xHA localization (red) in vacuoles undergoing bradyzoite differentiation following tachyzoite invasion (B) or in vacuoles following invasion with parasites further along the bradyzoite differentiation path (C). Bradyzoite differentiation was determined by SalmonE antibody to glycosylated CST1 (green in B), whereas anti-GFP antibody was used to identify differentiated bradyzoites expressing GFP under control of the bradyzoite specific LDH2 promoter (green in C). Nuclei labeled with DAPI. Scale bar equals 10 µm.

Applying the same Cas9 epitope tagging strategy used for MYR1 (Fig. 1A), the C-terminus of GRA16, GRA24, GRA28, and TgIST were epitope tagged separately in the PruΔku80 strain with the 3xHA tag. To assess effector export in these strains first under tachyzoite growth conditions, IFAs of fibroblast monolayers infected with tachyzoites of each effector-tagged strain were performed after fixing cultures in triplicate at 1, 2, and 3 days post-infection. In these experiments, effector intensity in host nuclei were exclusively measured in host cells containing single tachyzoite vacuoles at each timepoint. As reported previously, each of these effectors were detectable in host nuclei and were found to be significantly above baseline fluorescence levels (uninfected fibroblast nuclei) at 1 day p.i. (Fig. 2A-2D, asterisks). Although the nuclear intensities of all effectors were significantly above baseline out to three days post-infection, each effector demonstrated varying degrees of significant decline in host nuclei at later timepoints compared to their respective average values measured at 1 day p.i. (Fig. 2A-2D, hashtag symbols), indicating that each of these effectors do not continuously accumulate in the host nucleus during the course of tachyzoite infection.

**Figure 2.**
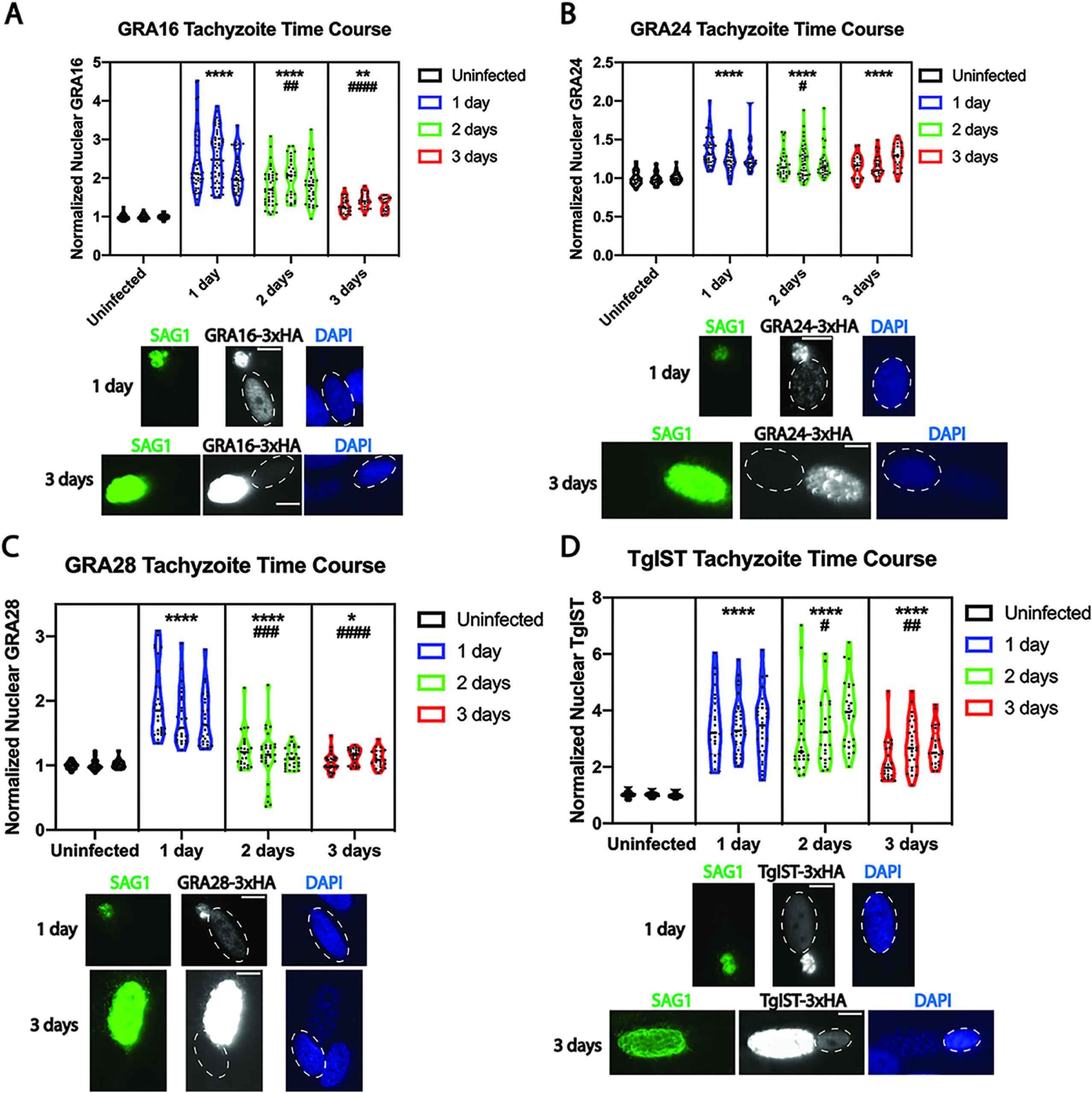
Quantitative analysis of exported effector fluorescent intensity in fibroblast host cell nuclei during tachyzoite infection reveal gradual declines in effector export over time. Violin plots and representative images of effector fluorescent intensity in infected fibroblast nuclei for GRA16-3xHA (A), GRA24-3xHA (B), GRA28-3xHA (C), and TgIST-3xHA (D) during the course of tachyzoite infection. Tachyzoite vacuoles were identified with antibody to SAG1 (green). Nuclei were labeled with DAPI (blue), and infected host cell nuclei are indicated by dashed ovals. Scale bar equals 10 µm. The mean gray values measured from host nuclei infected with single vacuoles were obtained from at least 15 fields of view and normalized to baseline fluorescence, measured from the nuclei of uninfected host cells. Three biological replicates are plotted for each time point, and mean gray values from at least 20 nuclei were measured from each replicate. Asterisks (*) indicate a statistically significant increase (**** p < 0.0001, *** p < 0.001, ** p < 0.01, * p< 0.05) compared to the uninfected group, whereas hashtag symbols (#) indicate a statistically significant decrease (#### p < 0.0001, ### p < 0.001, ## p < 0.01, # p < 0.05) in infected groups compared to the 1 day post-infection time point.

After confirming previously described tachyzoite effector export patterns, IFAs of the four tagged effector strains were performed under bradyzoite growth conditions, starting with egressed tachyzoites and inducing bradyzoite differentiation at the time of infection as described above. Monolayers were fixed at daily intervals from 1 to 6 days post-infection. The average fluorescent intensities of each effector were measured in the nuclei of host cells containing a single developing tissue cyst, ascertained either by positive BAG1 staining (1 to 3 days post-infection) or positive SRS9 staining (4 to 6 days post-infection), both markers of bradyzoite differentiation (19, 20) (Fig. 3A-D). All effectors were detectable in host nuclei at the earliest observed time point (1 day p.i.) and found to be significantly above average uninfected nuclear fluorescent intensity (Fig. 3A-D). Similar to measurements made from tachyzoite infected cells, the average host nuclear intensities of each effector significantly declined at later timepoints compared to intensities at 1 day p.i. (Fig. 3A-D). A divergence was noted in which effectors persisted in host nuclei at relatively late timepoints post-bradyzoite infection. Whereas TgIST and GRA16 remained significantly elevated compared to baseline in host nuclei out to 6 days p.i. (Fig. 3A, 3D), GRA24 and GRA28 levels were not significantly elevated starting at 2 days and 4 days p.i., respectively (Figure 3B, 3C). Despite their apparent absence in the host cell nucleus at later timepoints, GRA24 and GRA28 fluorescence was evident in differentiating bradyzoite vacuoles out to 6 days p.i. (Figure 3B, 3C, representative images), indicating that a secreted pool of GRA24 and GRA28 remains detectable in the vacuole, despite not being exported to the host cell nucleus, during the course of bradyzoite differentiation in this infection model.

**Figure 3.**
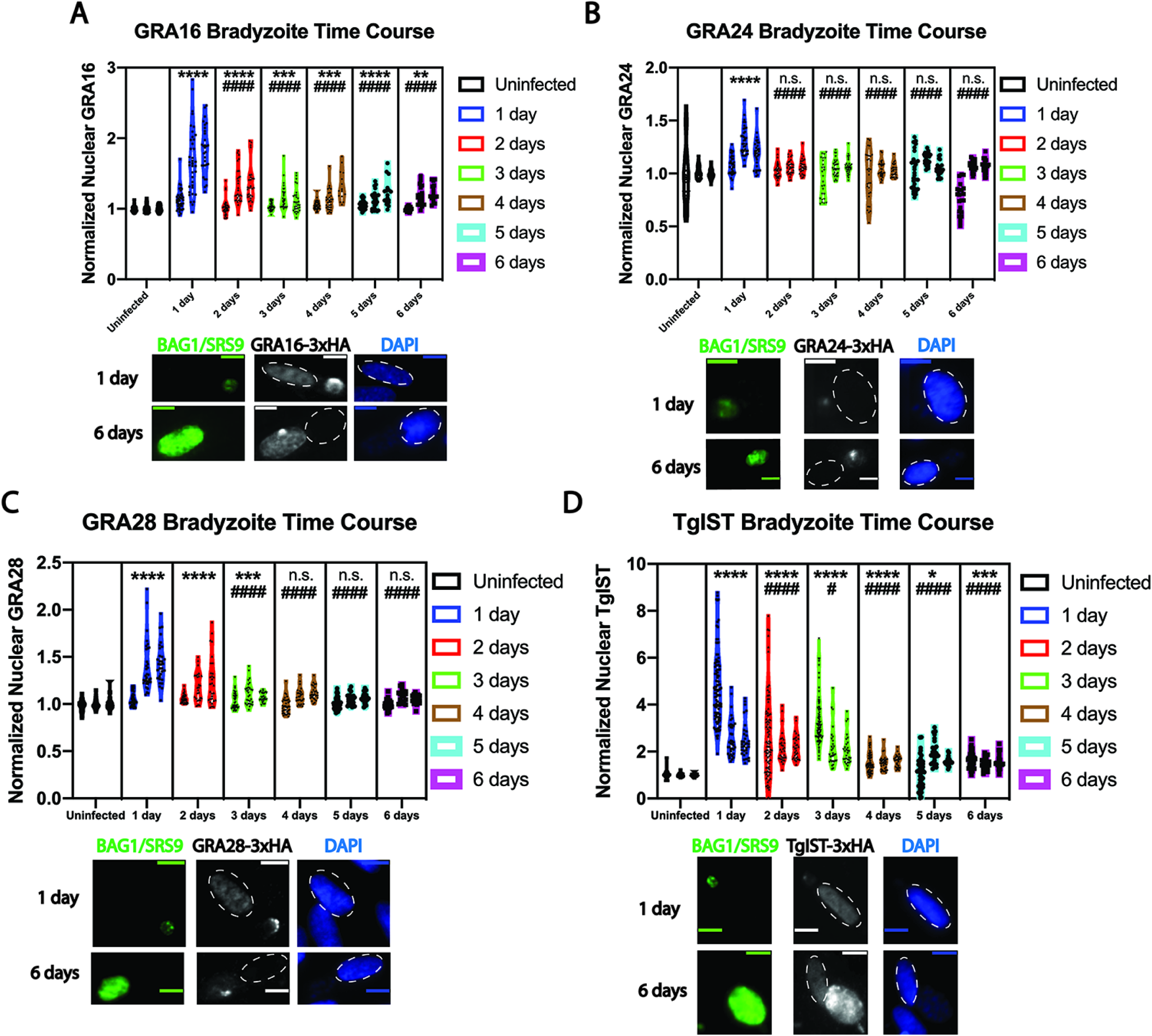
Quantitative analysis of exported effector fluorescent intensity in fibroblast host cell nuclei during bradyzoite differentiation reveal declines in effector export over time. Violin plots and representative images of effector fluorescent intensity in infected fibroblast nuclei for GRA16-3xHA (A), GRA24-3xHA (B), GRA28-3xHA (C), and TgIST-3xHA (D) during the course of bradyzoite infection. Bradyzoite differentiation was probed with antibody to either BAG1 at 1 day p.i. or to SRS9 at 6 days p.i. (both green). Nuclei were labeled with DAPI (blue), and infected host cell nuclei are indicated by dashed ovals. Scale bar equals 10 µm. Measurements and were made from three replicate experiments at each time point from infected nuclei containing a single differentiating vacuole at each timepoint, as described in Figure 2. The same mean gray value normalization approach and symbol conventions were used as in Figure 2 to indicate significant increases from the 1 day p.i. time point. Asterisks (*) indicate a statistically significant increase (**** p < 0.0001, *** p < 0.001, ** p < 0.01, * p< 0.05) compared to the uninfected group, whereas hashtag symbols (#) indicate a statistically significant decrease (#### p < 0.0001, ### p < 0.001, ## p < 0.01, # p < 0.05) in infected groups compared to the 1 day post-infection time point. n.s. indicates a non-significant difference compared to uninfected baseline values.

Quantitation of effector intensity in fibroblast nuclei at various timepoints demonstrated a decline as a function of time during tachyzoite to bradyzoite differentiation. We were interested in determining whether a similar pattern was also observed when infection was initiated by egressed bradyzoites harvested from *in vitro* bradyzoite-induced parasite cultures (as described above for MYR1 experiments). IFAs of HFFs infected with egressed *in vitro* induced bradyzoites revealed similar patterns of effector localization with respect to GRA16 and TgIST. GRA16 was detectable in the host nucleus at 1 and 2 days post-infection and appeared to decline in intensity at later timepoints (Fig. 4A), whereas TgIST was readily detectable at all time points (out to 4 days post-infection (Fig. 4D)). Intriguingly, GRA24 and GRA28 were not expressed at any time points using this infection model (Fig. 4B-C). The lack of expression of these two proteins in parasites farther along the bradyzoite differentiation path is supported by transcriptomic data deposited on ToxoDB (www.toxodb.org), in which several bradyzoite datasets demonstrate few to no transcripts from these two genes, suggesting that small amounts of these protein, if any, are translated in this life-stage. Hence, this finding demonstrates that the *in vitro* model of bradyzoite differentiation used here reflects data obtained from *in vivo* datasets.

**Figure 4.**
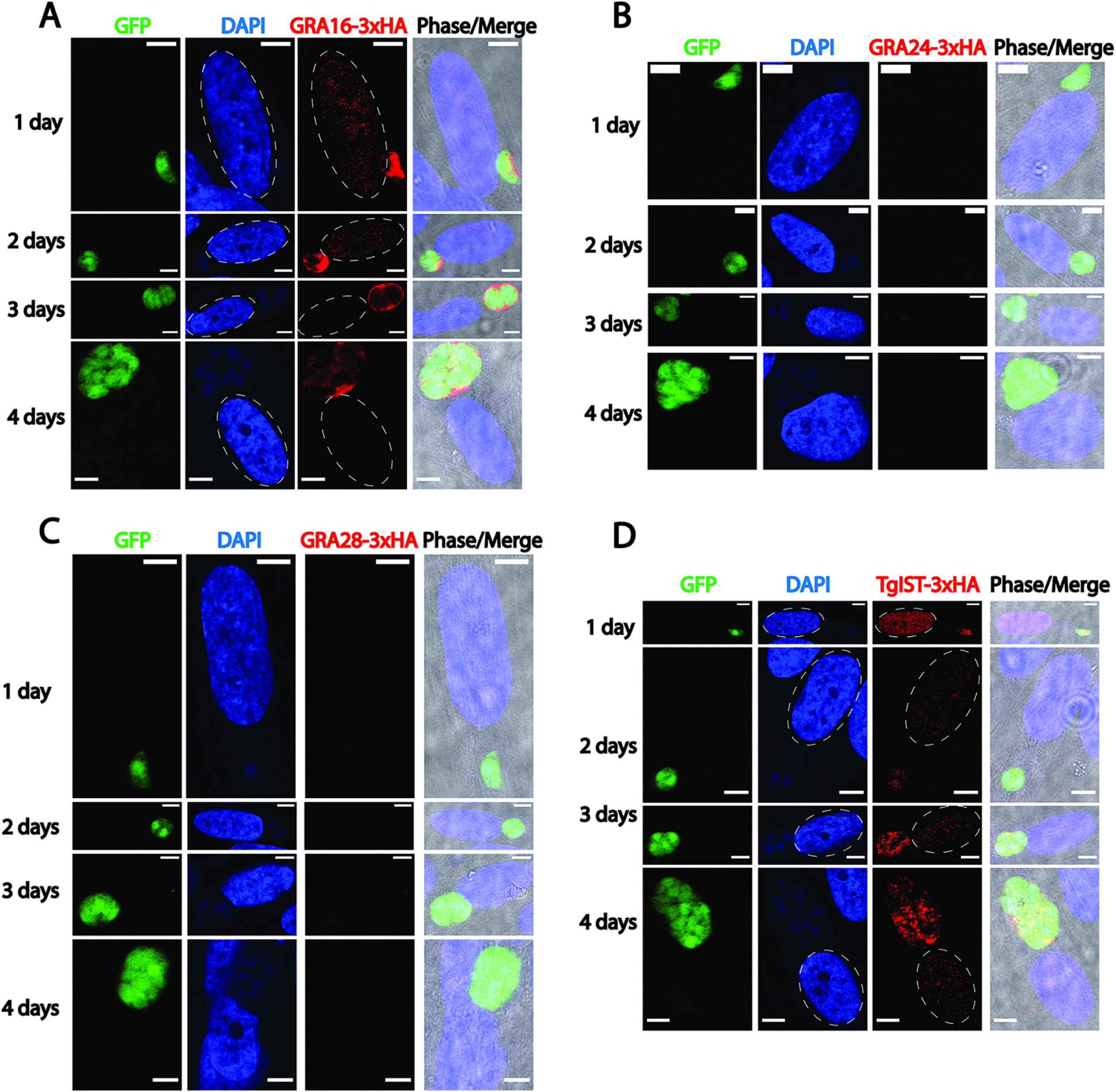
Immunofluorescence images of exported effector localization following infection with *in vitro* derived bradyzoites reveal differential expression of certain effectors. Representative images of GRA16-3xHA (A), GRA24-3xHA (B), GRA28-3xHA (C), and TgIST (D) localization (all in red) after infection of fibroblasts with egressed bradyzoites (e.g. parasites further along the bradyzoite differentiation path obtained *in vitro*, as determined by anti-GFP antibody [green]). Nuclei were labeled with DAPI (blue), and infected host cell nuclei are indicated by dashed ovals. Images were captured from fibroblasts infected with a single vacuole. GRA16 and TgIST are detected in infected host cell nuclei at 1 and 2 days post-infection, with TgIST readily detectable in host cell nuclei out to 4 days post-infection. GRA24 and GRA28 expression was undetectable. Scale bar equals 10 µm.

We were interested in measuring effector intensities in the nuclei of primary neurons, as neurons most frequently harbor tissue cysts in the mouse brain during chronic infection (21). Cortical neurons were harvested from mouse embryos after 14 days of gestation (E14) and were cultured *in vitro* for 14 days prior to infection (DIV14) on poly-L-lysine coated coverslips. Egressed tachyzoites from each effector-tagged strain were then used to infect separate neuron cultures, fixing them for IFAs at 1, 2, and 3 days post-infection. Staining with a monoclonal antibody to a bradyzoite secreted protein that we have named MAG2 (Gene ID: TGME49_209755; manuscript in revision), confirmed the occurrence of bradyzoite differentiation at 2 and 3 day p.i. (Fig. 5A-D). The nuclear effector intensities at 2 and 3 days p.i. were determined in neurons infected with a single MAG2 positive-vacuole, while nuclear intensities quantified at 1 day p.i. were obtained from infected neurons with single vacuoles as MAG2 expression was not observed at 1 day p.i. in neurons. Using ß-III tubulin as a neuron-specific marker, we found that host nuclear GRA24 was largely undetectable above baseline uninfected nuclear fluorescence at all timepoints, despite vacuolar expression (Fig. 5B). Of note, GRA24 export into neuron nuclei was detectable when the host neuron contained more than one vacuole at 1 day p.i. (data not shown). Similar to that observed in HFF host cells, GRA16 and GRA28 export was significantly elevated above baseline in neurons at 1 day p.i. and declined to baseline levels thereafter (Fig. 5A, 5C). TgIST remained significantly above baseline in host neuron nuclei at all timepoints inspected, displaying a trend of declining host nuclear intensity at 2 and 3 days p.i. (Fig. 5D). Thus, parasites retain the capacity to export each of these four effectors into the host nucleus during neuron infection, exhibiting a similar pattern of export to that observed in HFF host cells.

**Figure 5.**
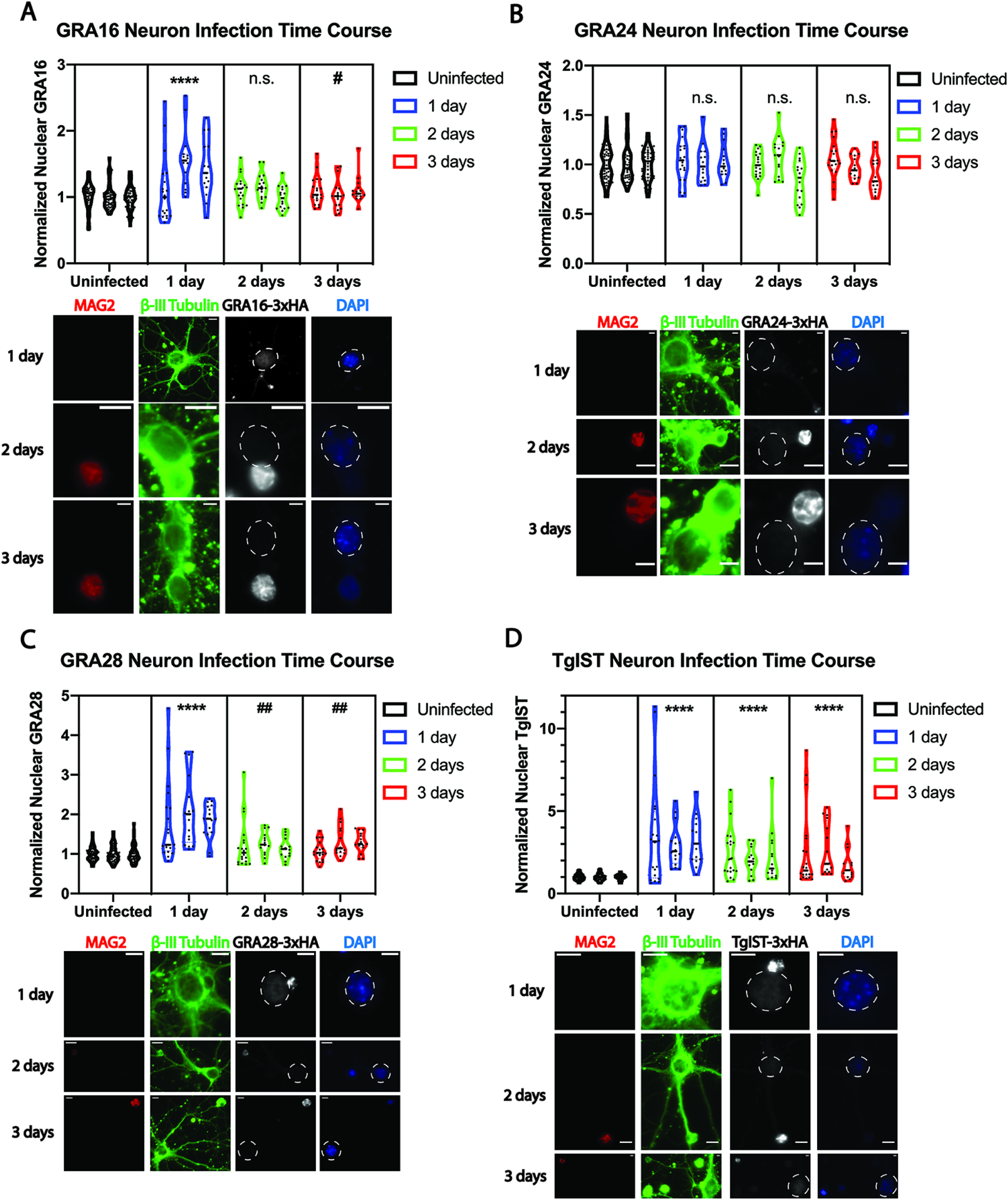
Quantitative analysis of exported effector fluorescent intensity in mouse primary cortical neuron nuclei reveal similar patterns of export compared to fibroblast infection. Violin plots and representative images of effector fluorescent intensity in infected primary neuron nuclei along with representative images at each timepoints for GRA16-3xHA **(A)**, GRA24-3xHA **(B)**, GRA28-3xHA **(C)**, and TgIST-3xHA **(D)** during the course of infection. Bradyzoite differentiation was determined with antibody to MAG2 (TGME49_209755) at 1, 2, and 3 days p.i. time points, though MAG2 positive vacuoles were not observed at the 1 day p.i. time point. Neurons were identified by positive ß-III tubulin staining (green). Nuclei were labeled with DAPI (blue), and infected host cell nuclei are indicated by dashed ovals. Scale bar equals 10 µm. Measurements and were made from three replicate experiments at each time point from infected nuclei containing a single vacuole at each timepoint. Mean gray values were normalized to uninfected nuclei, and symbol conventions were used as in Figure 2 to indicate significant increases from uninfected values or significant decreases from the 1 day p.i. time point. Asterisks (*) indicate a statistically significant increase (**** p < 0.0001, *** p < 0.001, ** p < 0.01, * p< 0.05) compared to the uninfected group, whereas hashtag symbols (#) indicate a statistically significant decrease (#### p < 0.0001, ### p < 0.001, ## p < 0.01, # p < 0.05) in infected groups compared to the 1 day post-infection time point. n.s. indicates a non-significant difference compared to uninfected baseline values.

The gradual decline of each exported effector observed during both tachyzoite and bradyzoite infection suggest that some factor(s) affecting effector export decreases in quantity or function during the course of vacuolar development. We sought to determine the mechanism behind this observation, reasoning that it could be due to a feature shared both by older bradyzoite and tachyzoite vacuoles. Recent work has demonstrated that catalytically active ROP17 is required for the translocation of protein effectors across the parasitophorous vacuole membrane (PVM) (22). ROP17, along with other rhoptry-derived proteins, is secreted at the time of host cell invasion, and ultimately localizes to the cytosolic face of the PVM. We speculated that declining levels of PVM-associated ROP17 during the course of intracellular infection may be the cause of declining effector translocation at later time points. To test this, we superinfected HFF monolayers containing parasites cultured under bradyzoite differentiation conditions for 7 days with a batch of egressed PruΔku80 strain tachyzoites that did not express the 3xHA-epitope tag. We allowed this superinfection to progress for one more day under bradyzoite differentiation conditions, and then performed IFAs on the monolayers, inspecting for a burst of host nuclear GRA16 or GRA28 in cells containing one differentiated vacuole (determined by fluorescent Dolichos lectin, DBA-FITC) and one new vacuole from superinfection. We found that no renewed GRA16 or GRA28 translocation was apparent under these conditions (Fig. 6A), even when HFFs contained multiple new vacuoles and a single bradyzoite vacuole, suggesting that the delivery of ROP17 from a new invasion event does not allow for the renewed export of vacuolar GRA16 and GRA28 from older vacuoles. The same results were obtained from an identical experiment performed under tachyzoite growth conditions, whereby tachyzoite infected host cells were superinfected at 2 days post-infection and fixed at 3 days post-infection (Fig. 6B). Previous groups have demonstrated that ROP17 provided by such superinfection reliablly restores the activity of ROP17 on the parasitophorous vacuole (22; Dr. J. Boothroyd personal communication). Thus, there are likely different mechanisms, either intra- or extra-vacuolar, that restrict translocation in older vacuoles.

**Figure 6.**
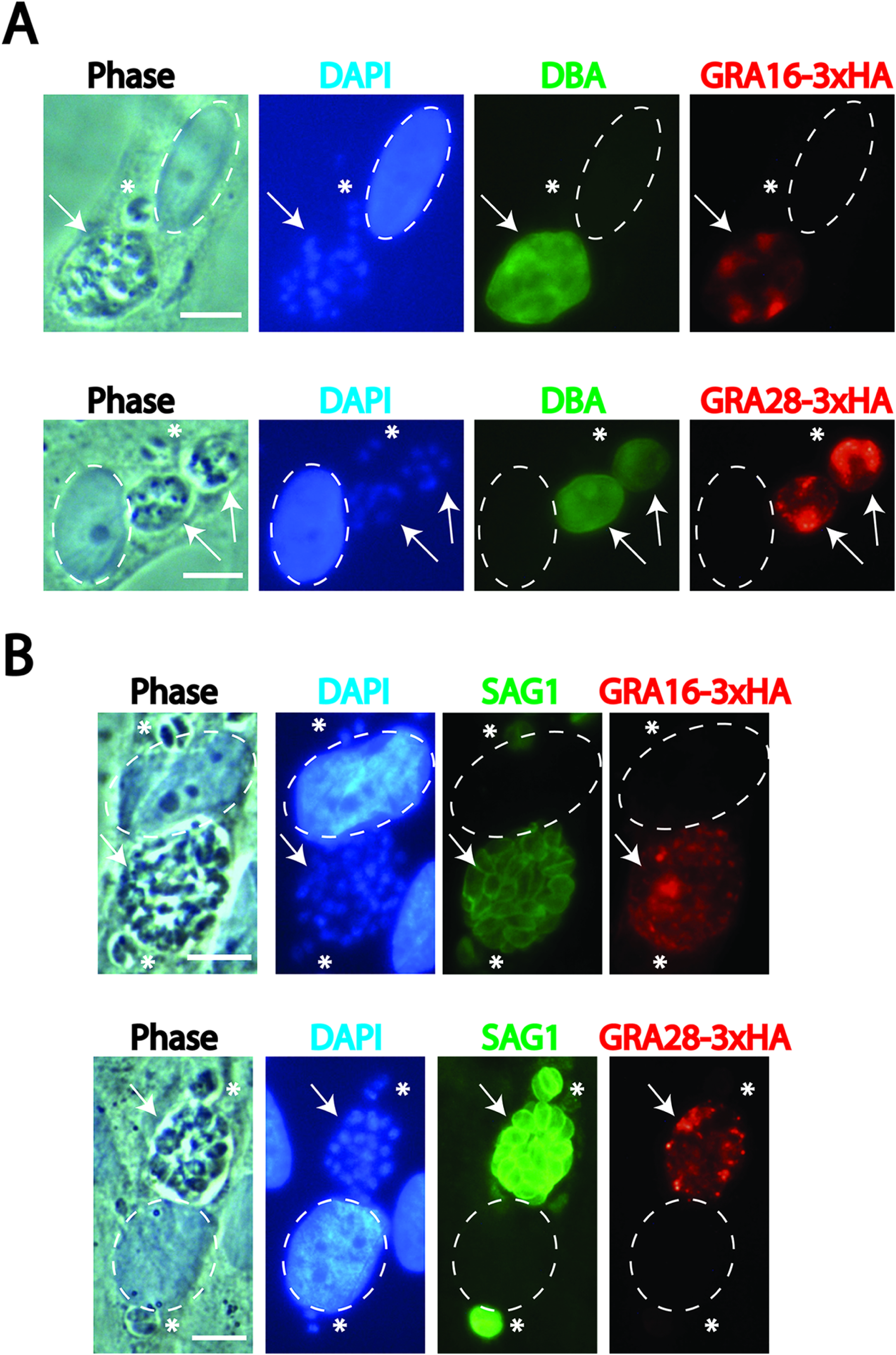
Immunofluorescence images of GRA16 and GRA28 localization following super-infection with non-epitope-tagged *T. gondii* tachyzoites demonstrate no renewed effector export. **(A)** Representative images of GRA16-3xHA and GRA28-3xHA localization (both in red) 1 day post-tachyzoite invasion and 8 days post-differentiation (tachyzoites introduced 7 days post-differentiation). Differentiated vacuoles are indicated by DBA-FITC staining (green). Nuclei were labeled with DAPI (blue), and infected host cell nuclei are indicated by dashed ovals. Arrows indicate the bradyzoite containing vacuoles with vacuolar GRA16 or GRA28, whereas asterisks indicate tachyzoite vacuoles in the same host cell. Neither GRA16 nor GRA28 are exported from differentiated vacuoles into the host cell nucleus 1 day post-tachyzoite invasion. Scale bar equals 10 µm. **(B)** Representative images of GRA16-3xHA and GRA28-3xHA localization (both in red) 3 days post-initial infection and 1 day post-tachyzoite superinfection (tachyzoites introduced 2 days post-infection). Older vacuoles (arrow) from the initial infection are larger than vacuoles from superinfection (asterisk), though both young and older tachyzoite vacuoles are labeled by SAG1 (green). Nuclei were labeled with DAPI (blue), and infected host cell nuclei are indicated by dashed ovals. Scale bar equals 10 µm.

Robust TgIST export is notable both during tachyzoite infection (Fig. 2D) and during the bradyzoite infection models used in this study (Fig. 3D, 4D). TgIST has been shown to suppress the host cell response to IFN-γ, preventing the upregulation of genes downstream of this signaling pathway, such as interferon response factor 1 (IRF1). Given the known function of TgIST and the findings obtained by IFA, we hypothesized that cells infected with differentiating bradyzoites remain unresponsive to exogenous IFN-γ at relatively late timepoints post-infection. Using IRF1 as a probe for IFN-γ signaling, we measured host nuclear IRF1 fluorescence intensity in HFFs harboring a single tissue cyst and stimulated with recombinant IFN-γ 7 days post-infection. The results from multiple IFAs demonstrated that a significant attenuation of IRF1 upregulation occurred in HFFs infected with parasites of the PruΔku80 strain, compared to uninfected cells and cells infected with TgIST-KO parasites (Fig. 7A-B). Hence, in the context of prolonged bradyzoite infection, TgIST appears to be at least partially responsible for suppressing IFN-γ signaling in the host cell.

**Figure 7.**
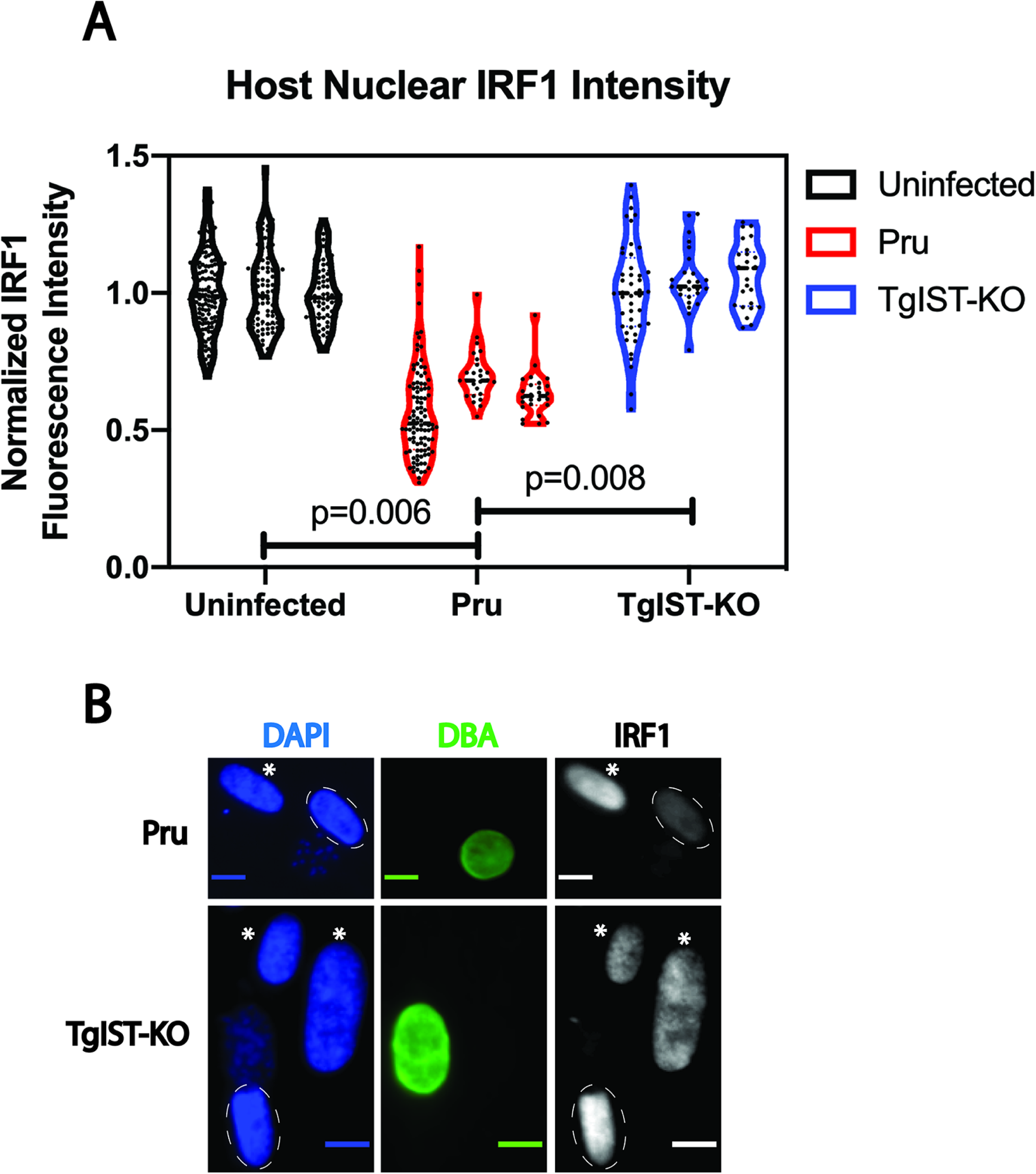
Quantitation and immunofluorescence images of IRF1 upregulation in IFN-γ stimulated fibroblasts infected with either Pru or TgIST-KO demonstrate that IRF1 suppression is TgIST dependent during bradyzoite infection. **(A)** Violin plots of IRF1 intensity in the nuclei of uninfected fibroblasts or infected fibroblasts containing either PruΔku80 or Pru TgIST-KO parasites, measured from three replicates. The mean fluorescent intensity of IRF1 was normalized to the average values obtained from uninfected nuclei. The IRF1 intensity values plotted for the uninfected group were obtained from both PruΔku80 and Pru TgIST-KO infected cultures for each replicate. **(B)** Representative images of fibroblasts infected initially with egressed tachyzoites from the PruΔku80 strain or Pru TgIST-KO strain, 7 days post-infection in bradyzoite inducing conditions. Fibroblasts were exposed to IFN-γ at a concentration of 100U/mL for 6 hours at 7 days post-infection. IFN-γ signaling was probed with anti-IRF1 antibody (red), and bradyzoite differentiation was determined by DBA-FITC staining (green), a cyst wall marker. Nuclei were labeled with DAPI (blue), and images were captured from fibroblasts infected with a single vacuole. Notable IRF1 expression is observed in uninfected nuclei (asterisks), whereas IRF1 expression is attenuated in infected host cell nuclei (dashed ovals) containing Pru parasites, but not TgIST-KO parasites. Scale bar equals 10 µm.

The notable persistence of TgIST in the host nucleus at 6 days post-infection, and the prolonged effects mediated by this protein in the host cell, raised the possibility that TgIST could be an effector that is continuously exported, albeit at a reduced efficiency during late infection. Alternatively, TgIST persistence at late time points could be due to a relatively large bolus of TgIST export during the early phase of infection. To address these possibilities, we sought to determine the stability of TgIST in the host nucleus using a previously described inducible knockout strategy, in which genetic deletion is achieved using exogenous rapamycin to dimerize Cre fragments and induce Cre enzymatic activity (23). A single plasmid containing the following was constructed after several cloning steps: floxed TgIST-3xHA tagged locus driven by the endogenous TgIST promoter and TgIST 5’UTR, YFP-3xMyc tagged reporter protein (the expression of which is induced only after Cre-mediated removal of the TgIST coding sequence) and open reading frames encoding two “DiCre” fragments (Cre59 and Cre60), among other elements (Fig. 8A). PruΔku80 strain tachyzoites were transfected with the afore-mentioned linearized construct (TgIST-GeneSwap-DiCre). PCR of genomic DNA from subcloned parasites revealed that the construct had inserted as a second copy into the parasite genome (data not shown), therefore, in these subsequent experiments with the TgIST-GeneSwap-DiCre parasites, the ectopic 3xHA-epitope tagged copy of TgIST driven by its own endogenous promoter was detected by IFA and then deleted by DiCre. To evaluate the DiCre parasites, cultures infected for 1 day under bradyzoite growth conditions were exposed to 50nM rapamycin for 24hrs, after which rapamycin containing media was removed and replaced with fresh media. IFAs of these cultures at 5 days p.i. revealed that the vast majority of TgIST-GeneSwap-DiCre parasites expressed the YFP-3xMyc reporter protein and only trace amounts of TgIST-3xHA, whereas in the absence of rapamycin exposure, most TgIST-GeneSwap-DiCre parasites expressed robust TgIST-3xHA expression and no YFP-3xMyc reporter protein (Fig. 8C, representative images).

**Figure 8.**
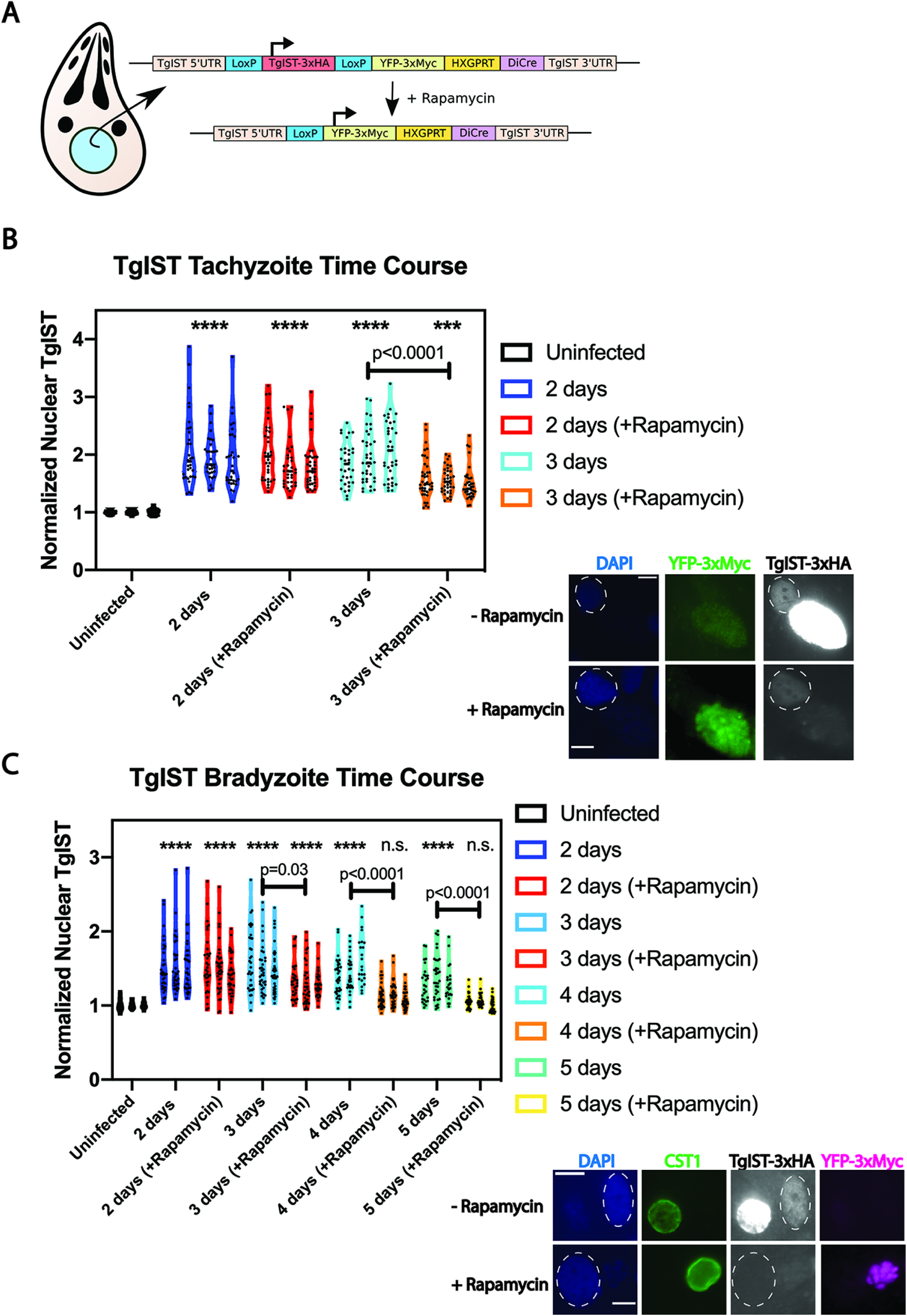
TgIST-GeneSwap-DiCre parasites allow for conditional knockout of ectopic TgIST-3xHA, revealing that stability of TgIST in host fibroblast nuclei is limited to several days during the course of bradyzoite differentiation. **(A)** Illustration of the inducible TgIST-3xHA knockout strategy. Upon the addition of rapamycin, the TgIST-3xHA transgene is deleted, allowing for the expression of YFP-3xMyc driven by the TgIST promoter and 5’UTR sequence. Both constitutively expressed DiCre fragments and an HXGPRT selectable marker are included in the same construct. **(B)** Violin plots of normalized TgIST fluorescent intensity in host fibroblast nuclei during the course of tachyzoite infection. Measurements were made from three replicate experiments at each time point from infected nuclei containing a single vacuole at each timepoint. Mean gray values were normalized to uninfected nuclei. Symbol conventions were used as in Figure 2 to indicate significant increases from uninfected values (asterisks). A significant decrease in normalized TgIST intensity was observed in rapamycin treated coverslips compared to untreated coverslips at 3 days p.i. Representative images of TgIST-GeneSwap-DiCre parasites 3 days post-infection with or without a 24hr 50nM rapamycin pulse exposure 1day post-infection, allowing for an initial TgIST secretion event. Nuclei are labeled with DAPI (blue), YFP-3xMyc reporter with Myc-tag antibody (green), and TgIST-3xHA labeled with HA antibody. Infected host cell nuclei are indicated by dashed ovals. Scale bar equals 10 µm. **(C)** Violin plots of normalized TgIST fluorescent intensity in host fibroblast nuclei during the course of bradyzoite infection. Measurements were made from three replicate experiments at each time point from infected nuclei containing a single vacuole at each timepoint. Mean gray values were normalized to uninfected nuclei. Symbol conventions were used as in Figure 2 to indicate significant increases from uninfected values (asterisks). A significant decrease in normalized TgIST intensity was observed in rapamycin treated coverslips compared to untreated coverslips at 3, 4, and 5 days p.i. Host nuclear TgIST was not found to be significantly elevated above uninfected values in rapamycin treated coverslips at 4 and 5 days p.i. (n.s.). Representative images of TgIST-GeneSwap-DiCre parasites 5 days post-bradyzoite differentiation with or without a 24hr 50nM rapamycin pulse exposure 1day post-infection. Bradyzoite differentiation was assessed by SalmonE antibody to glycosylated CST1 (green), nuclei labeled with DAPI (blue), TgIST-3xHA labeled with HA antibody, and the YFP-3xMyc reporter with Myc-tag antibody (magenta). Infected host cell nuclei are indicated by dashed ovals. Scale bar equals 10 µm.

A time course was obtained for ectopic TgIST-3xHA expression in the TgIST-GeneSwap-DiCre parasites in both tachyzoites and bradyzoite. TgIST-GeneSwap-DiCre parasites were allowed to infect HFF monolayers under tachyzoite or bradyzoite growth conditions and a rapamycin pulse was provided at 1 day p.i. IFAs were performed daily following the rapamycin pulse to determine how host nuclear TgIST intensity changed, comparing rapamycin treated and untreated TgIST-GeneSwap-DiCre parasites. In these experiments, only host cells containing YFP-3xMyc positive parasites cells were analyzed in the rapamycin treated group, whereas in the untreated group, only host cells containing YFP-3xMyc negative parasites were analyzed. Under tachyzoite growth conditions, a significant decrease in normalized nuclear TgIST was evident in host cells from rapamycin treated coverslips compared to untreated cultures at 3 days post-infection (Fig. 8C); however, at all timepoints, TgIST was detectable above baseline fluorescence even after TgIST deletion at 1 day p.i. (Fig. 8C, asterisks), indicating at least some portion of TgIST protein exported during the first day of infection persists in the host nucleus after three days of tachyzoite infection. Intriguingly, we found a similar decrease in TgIST nuclear intensity during the bradyzoite time course experiment at 3 days p.i., although at 4 and 5 days p.i., host nuclear TgIST was no longer detectable above baseline fluorescence (Fig. 8D). Rapamycin pulse exposure itself does not affect TgIST-3xHA stability in host nuclei, as experiments with endogenously tagged TgIST-3xHA tagged parasites without the DiCre transgene demonstrated TgIST host nuclear persistence above baseline at 3, 4, and 5 days p.i. when exposed to a 24hr rapamycin pulse at 1 day p.i. (Fig. S2). Hence, genetic deletion of ectopic TgIST on day 1 post-infection indicates that this protein is exported beyond day 1, and that following export on day 1 TgIST can persist at some level in the host nucleus out to about 4 days post-infection.

## DISCUSSION

Our findings demonstrate that parasite protein translocation across the nascent cyst membrane is a feature of bradyzoite infection both in fibroblasts and neurons. We set about to test our initial hypothesis of continuous bradyzoite effector export after our previous finding of MYR1 as a putative cyst wall protein (6). The expression of MYR1 within *in vitro* tissue cysts at various timepoints was validated, as this protein is secreted into differentiating vacuoles regardless of whether the invading parasite more closely resembled a tachyzoite or bradyzoite (Fig. 1 and Fig. S1). The presence of MYR1 in differentiating tissue cysts clearly allows for effector export to occur at the early stages of infection, though seemingly not at later stages of infection in the case of GRA24 and GRA28 (Fig. 3B-C). The differences in host nuclear intensities between GRA16/TgIST and GRA24/GRA28 at late timepoints may reflect decreased transcription of GRA24 and GRA28 during bradyzoite differentiation, as documented by various bradyzoite datasets on ToxoDB. Intriguingly, we found that invasion by *in vitro* derived egressed bradyzoites did not result in the export of either GRA24 or GRA28, as neither protein was expressed during this infection model (Fig. 4). This finding suggests that, depending on where the parasite lies on the bradyzoite differentiation continuum, a different arsenal of exported effectors may be utilized by the bradyzoite. This could allow for the parasite to refine the manner of host cell manipulation. For example, a previous report has linked GRA24 as a negative regulator of bradyzoite differentiation (24). In this context, GRA24 export from a differentiating vacuole could interfere with the process of bradyzoite and cyst maturation.

The export of all four effectors studied here could be detected in mouse primary cortical neurons, though GRA24 export was undetectable when analyses were limited to single vacuole infections (Fig. 5B). This finding suggests that effector export may occur in this cell type during the establishment of chronic infection *in vivo*, though clearly further investigation is needed to determine the nature of parasite protein export from the parasitophorous vacuole *in vivo* during acute and chronic infection in sites such as skeletal muscle and brain. We found that while similar patterns of export were identified in neurons compared to fibroblast infection (robust export at 1 day p.i., followed by decline thereafter), the decline to baseline host nuclear fluorescence levels was more rapid in neurons for both GRA16 and GRA28. As each of these effectors were detectable in the parasitophorous vacuole during neuron infection, we reason that this finding could reflect cell-type specific differences in nuclear effector stability, or that effector export is less efficient in neurons. These findings may partially explain the previously reported cell-type dependent transcriptional responses to *T. gondii* in fibroblasts, neurons, astrocytes, and skeletal muscle cells (25).

It is apparent that all effectors accumulate within developing tissue cysts *in vitro* during the tachyzoite-to-bradyzoite transition, possibly due to a decreased capacity or efficiency to export these proteins. This finding could reflect the maturation of the cyst wall and function of the cyst wall as a barrier to large molecule transport, as previously reported using fluorescent dyes of various sizes (26). On the other hand, we do not exclude the possibility that these effectors may perform additional functions within the tissue cyst. The mechanism behind the decline in effector export at later timepoints is unclear, and may not occur uniquely in bradyzoite vacuoles, as a time course of tachyzoite infection reveals a similar pattern (Fig. 2). None of these protein effectors appeared to be differentially processed at late time points post-bradyzoite differentiation when protein migration was assessed by immunoblotting (data not shown).

In light of the recently described role of the rhoptry protein ROP17 on the export of MYR-dependent effectors (27), we tested whether new infections with egressed tachyzoites (providing fresh ROP17 in-trans) could initiate the translocation of effectors present in older tachyzoite or bradyzoite vacuoles and found that no renewed export was detected (Fig. 6). Hence, declining ROP17 levels during infection does not appear to be the cause of export decline. In any case, the pattern of protein export observed for the effectors in this study suggest a common need shared by both tachyzoite and bradyzoites in rewiring the host cell during the early stages of infection, inducing changes that may persist long after the degradation of these effectors in the host cell.

Despite an apparent decline in effector export, we were interested in determining how long TgIST persisted in fibroblast nuclei, and by extension whether TgIST might be a continuously exported effector. We engineered a parasite strain expressing a floxed ectopic epitope tagged copy of TgIST that could be deleted by inducible Cre excision, and found that when deleting this copy at 1 day p.i., TgIST was no longer detectable during bradyzoite differentiation at 4 and 5 days p.i. (Fig. 8D). We reason that if a bolus of TgIST exported during the first day of infection persists in host fibroblast nuclei for only 4 days, and if TgIST export can be typically detected above baseline out to 7 days p.i., then there are export events occurring at least at 2 or 3 days post-infection. Further experiments are needed to model TgIST export during bradyzoite infection in more detail. If TgIST export occurs at a basal level at late timepoints, this may position TgIST as a unique exported effector among the four studied here, as a previous study demonstrated the absence of GRA16 and GRA24 export into the host cell when expression was initiated 5 days post-bradyzoite differentiation using a Tet operator approach (28).

The prolonged suppression of IFN-γ signaling in bradyzoite infected fibroblasts is not surprising, given the persistence of TgIST in the host cell nucleus during bradyzoite infection compared to other effectors. TgIST has been shown to simultaneously bind STAT1 and components of the Mi-2/NuRD protein complex, leading to altered chromatin marks at genomic loci recognized by STAT1 and preventing STAT1-mediated transcription (11, 12). TgIST-dependent attenuation of IFN-γ signaling was demonstrated in bradyzoite infected fibroblasts (Fig. 7), suggesting that despite declining levels of TgIST at later time points post-infection, the altered chromatin state at STAT1-binding loci may be responsible for long-lasting STAT1 transcriptional repression in the infected host cell. As no robust increase of nuclear IRF1 was detected in neurons following IFN-γ exposure of various lengths (data not shown), a similar attenuation in IRF1 due to TgIST could not be demonstrated in primary cortical neurons. IFN-γ signaling in neurons is currently not well characterized, although reports in primary mouse hippocampal neuron cultures demonstrate an intact albeit delayed IFN-γ signaling pathway (29) (30). Indeed, this delayed neuronal signaling cascade downstream to IFN-γ may make this cell type more vulnerable to successful parasite infection and could partially explain why tissue cysts are most frequently found in neurons in the brain.

GRA16 export occurs early following *in vitro* derived bradyzoite invasion of a new host cell. GRA16 has been shown to bind to HAUSP and PP2A in various human and mouse cell lines, and substantial evidence has been provided to show this protein is likely involved in the G2/M phase arrest of the infected host cell (8). It is known that *in vivo* bradyzoite cysts typically reside in neurons in the brain and skeletal muscle cells in muscle tissue during chronic infection (21) (31), both cell types that are terminally differentiated with respect to their cell cycle. We speculate that GRA16 may play a more relevant role in bradyzoite infection of other cell types (i.e. intestinal epithelial cells) following carnivory of tissue containing tissue cysts, where host cell cycle control is likely more important for successful replication and dissemination in a new host. However, one cannot exclude the possibility that GRA16 serves additional functions in terminally differentiated cell types, or potentially interacts with other host proteins in different cell types.

In summary, the data presented here extend our knowledge of an intriguing method used by this parasite to manipulate the cells it infects, and points toward one mechanism responsible for the persistence of this organism during chronic infection of its host.

## MATERIALS AND METHODS

### Cell culture

PruΔku80Δhxgprt LDH2-GFP parasites (18) were continuously passaged in human foreskin fibroblast (HFF : ATCC:CRL-1634; Hs27) host cells, as previously described (32). TgIST-KO parasites were obtained as a kind gift from the laboratory of Dr. David Sibley (Washington University School of Medicine, St. Louis, MO). For all experiments using fibroblasts, bradyzoite induction was performed at the time of invasion by replacing growth media with bradyzoite induction media (50 mM Hepes, pH8.2, DMEM supplemented with 1% FBS, penicillin and streptomycin) prior to infection with egressed tachyzoites at a multiplicity of infection (MOI) of 1 when infecting confluent HFF monolayers on glass coverslips or at a MOI of 2 in T25 flasks. Bradyzoite induction cultures were incubated in a humid 37 °C incubator without CO_2_. For infection of coverslips, induction medium was changed every 2 days. To obtain egressed bradyzoites from T25 flasks, induction media was changed only on the second day of infection. Parasites i.e. egressed bradyzoites were harvested from cultures 6 days post-induction, when over 95% of *T. gondii* demonstrated GFP expression in the culture.

Mouse primary cortical neurons were harvested from E14 mouse embryos obtained from pregnant C57Bl/6 mice, ordered from Charles River. Dissections of E14 cortical neurons were performed as previously described (33). Following dissection, 30,400 cortical neurons were added to poly-L-lysine coated glass coverslips in 24-well plates and later cultured in Neurobasal Media (Thermo Fisher) supplemented with GlutaMAX Supplement (Thermo Fisher) and B-27 Supplement (Gibco). After 4 DIV, cytarabine (ara-C) was added to each culture at a final concentration of 0.2µM to minimize contamination from dividing, non-neuronal cells. Cultures were maintained for up to 18 days by replacing half of the conditioned media with fresh supplemented Neurobasal media every 7 days.

For TgIST-DiCre timecourse experiments, TgIST-GeneSwap-DiCre parasites (or control endogenously tagged TgIST-3xHA parasites without the DiCre transgene) were allowed to infect HFF monolayers under tachyzoite growth conditions or induced to differentiate into bradyzoites as described above. At 24hrs post-infection, tachyzoite or bradyzoite differentiation media was replaced with fresh media supplemented with 50nM rapamycin. After 24hrs of rapamycin exposure, cell media was replaced with fresh media without additional rapamycin, and infected cells were cultured in the absence of exogenous rapamycin thereafter.

### Cloning and parasite transfection

To epitope tag the genes described in this study at their endogenous loci using CRISPR/Cas9, single guide RNAs (sgRNA) targeting the C-terminus or 3’UTR of each gene of interest were cloned separately into the p-HXGPRT-Cas9-GFP plasmid backbone using KLD reactions, as previously described (34). Donor sequences for homology mediated recombination were generated by amplifying a 3xHA tag, the 3’UTR of HXGPRT, and a DHFR mini-cassette to confer pyrimethamine resistance from the previously described pLIC-3HA-DHFR plasmid backbone (6) (Fig. 1A). Primers used to amplify this donor sequence also contained overhangs with at least 30bp homology to the C-terminus or 3’UTR of the gene of interest.

To engineer a floxed TgIST-3xHA vector that also contained a YFP-3xMyc reporter gene, a HXGPRT selectable marker, and the Cre59 and Cre60 DiCre fragments under control of robust eukaryotic promoters on a single vector (TgIST-GeneSwap-DiCre), several sequential Gibson Assembly reactions were performed. Briefly, in the first assembly, a YFP open reading frame (ORF) and a HXGPRT selectable marker were PCR amplified from a GeneSwap plasmid backbone (kind gift from Dr. Kami Kim) and concatenated using a forward primer containing a LoxP sequence and homology to the HXGPRT 3’UTR sequence downstream of a pLIC-TgIST-3xHA construct containing the endogenous TgIST promoter and 5’UTR, and a reverse primer containing homology to a DiCre plasmid backbone containing the promoters, 3’UTRs, and ORFs of the Cre59 and Cre60 DiCre fragments (kind gift from Dr. Kami Kim). In the second assembly, the concatenated construct from the first assembly was PCR amplified and further concatenated to include a LoxP sequence upstream of the TgIST start codon, as well as a homologous arm to the endogenous 3’UTR of TgIST downstream of the DiCre sequences. A KLD reaction was performed to introduce a 3xMyc epitope tag at the C-terminus of the YFP ORF using the second Gibson Assembly reaction product. The final Gibson Assembly was performed to re-introduce the Cre59 promoter, ORF, and 3’UTR from the DiCre plasmid backbone after it was found to be absent in the second Gibson Assembly product. A plasmid map of the final TgIST-GeneSwap-DiCre construct is available on request. A full list of primers used for cloning and epitope tagging can be found in Supplemental Table S1.

Pru*Δku80Δhxgprt* tachyzoites were transfected with 7.5 μg Cas9 plasmid and 1.5ug PCR amplified donor sequence with electroporation in cytomix. Transfected parasites were selected for with 2μM pyrimethamine for at least two passages, after which resistant parasites were subcloned by limiting dilution. For transfection of Pru*Δku80Δhxgprt* tachyzoites with the TgIST-GeneSwap-DiCre vector, 7.5 μg Cas9 plasmid targeting the TgIST C-terminus was co-transfected with 40ug of the TgIST-GeneSwap-DiCre vector linearized with ScaI restriction enzyme (NEB). Drug selection with DMEM complete media containing 25µg/mL mycophenolic acid and 50µg/mL xanthine was performed 24hrs post-transfection for the following 6 days before removing selection media and subcloning by limiting dilution after sufficient parasite egress was observed.

### Immunofluorescence Assays

Confluent HFF monolayers and primary neurons were cultured on glass coverslips and infected with either egressed tachyzoites or egressed bradyzoites differentiated *in vitro* following 6 days of growth under bradyzoite inducing conditions. All infections occurred under bradyzoite inducing conditions at the time of infection. For the IFN-γ stimulation experiments, HFF monolayers infected initially with tachyzoites and induced to differentiate into bradyzoites were stimulated with 100U/mL IFN-γ 7 days post-infection and fixed 6 hours post-stimulation. All coverslips were fixed with 4% PFA for 20 minutes at room temperature, permeabilized in a 0.2% Triton X-100, 0.1% glycine solution for 20 minutes at room temperature, rinsed with PBS, and blocked in 1% BSA for either 1 hour at room temperature or at 4C overnight. Coverslips were next labeled as follows: HA-tagged proteins were detected by anti-HA rat monoclonal antibody 3F10 (Sigma, 1:200), parasite cyst wall and cyst matrix by in-house SalmonE anti-CST1 antibody (1:500) and in-house anti-MAG2 antibody (1:500) respectively, bradyzoites by anti-BAG1 (1:500) and rabbit anti-SRS9 (1:1000), GFP by rabbit anti-GFP (Thermo Fisher, 1:500), IRF1 by rabbit anti-IRF1 (Cell Signaling, 1:500), tachyzoite SAG1 by anti-SAG1 (Thermo Fisher, 1:500), and neurons by anti-ß-III Tubulin (1:1000, kind gift from Dr. David Sharp, Albert Einstein College of Medicine, Bronx, NY). Rabbit anti-Myc tag antibody (Cell Signaling Technologies, 1:500) was used to detect YFP-3xMyc reporter expression in TgIST-DiCre infected coverslips. Various combinations of Alexa 488 conjugated anti-rabbit and anti-mouse, Alexa555 and Alexa594 conjugated anti-rat, and Alexa633 conjugated anti-rabbit antibodies were used as secondary antibodies at a dilution of 1:1000 (Thermo Fisher). DAPI counterstain was used to label parasite and host cell nuclei (1:2000). Coverslips were mounted in ProLong Gold Anti-Fade Reagent (Thermo) and imaged using either a Leica SP8 confocal microscope, a Nikon Eclipse widefield fluorescent microscope (Diaphot-300), or a Pannoramic 250 Flash III Automated Slide Scanner (3D Histech).

### Quantitative Image Analysis

For each replicate and timepoint, and for every time course, at least 15 randomly selected fields of view were either obtained and exported from CaseViewer (3D Histech) software after acquiring images with a Pannoramic 250 Automated Slide Scanner or obtained from imaging with a Lecia SP8 confocal microscope. For each effector time course, identical exposure times were used to detect effector fluorescence intensity from each coverslip. Images obtained from either the confocal or slide scanner microscopes were viewed in ImageJ (NIH), and nuclei were segmented to generate regions of interest. Regions of interest (nuclei) were selected manually from fibroblasts or neurons that contained individual parasitophorous vacuoles, and the mean gray value was measured from each region of interest in the fluorescent channel used to detect each effector. Background fluorescence was determined by selecting nuclei from fibroblasts or neurons that did not contain parasitophorous vacuoles (uninfected), measuring mean gray values as described above. To allow for comparisons between replicate experiments and across various timepoints, mean gray values measured from infected host nuclei were normalized by dividing each measured value by the average background fluorescence quantified from uninfected nuclei at each timepoint (i.e. from each coverslip). Normalized values were plotted using PRISM 8 (GraphPad). As each normalized mean gray value dataset was shown not to exhibit a Gaussian distribution after testing for normality, the nonparametric Kruskal-Wallis test along with Dunn’s multiple comparisons test was used to compare the means from three replicates for a given timepoint to determine statistically significant differences between groups.

## ACKNOWLEDGEMENTS

We thank members of the Weiss lab for their comments, suggestions, and insights in the preparation of this manuscript. We thank Drs. John Boothroyd, Jeroen Saeij, and David Sibley for useful discussions and suggestions on the experiments performed in this study. We thank the Albert Einstein Analytical Imaging Facility, specifically Dr. Vera DesMarais and Andrea Briceno for training on various microscopes and suggestions for ImageJ analysis.

This work was supported by P30CA013330, SIG #1S10OD016214-01A1, and SIG #1S10OD019961-01 (Einstein Analytical Imaging Facility), 1F31AI136401 (J.M.), R01AI134753 (L.M.W.), and R21AI123495 (L.M.W.).

## Author contributions

J.M. and L.M.W. conceived and designed the work; J.M. and P.S. performed the experiments and analyzed data obtained from experiments; J.M. and L.M.W. wrote the paper.

## SUPPLEMENTAL FIGURES AND TABLES

**Supplementary Table S1.**

List of primers used in this study for CRISPR/Cas9 tagging of genes and for TgIST-GeneSwap-DiCre construction.

**Figure S1. Immunofluorescence images of MYR1-3xHA tagged parasites during bradyzoite differentiation 2-6 days post-infection.**

Representative images of MYR1-3xHA localization (red) in vacuoles undergoing bradyzoite differentiation, as determined by SalmonE antibody to CST1 (green). Nuclei labeled with DAPI. Scale bar equals 10 µm.

**Figure S2. Rapamycin pulse exposure does not alter the observed stability of TgIST-3xHA in host fibroblast nuclei**

**(A)** Violin plots of normalized TgIST fluorescent intensity in host fibroblast nuclei during the course of bradyzoite infection with the endogenously tagged PruΔku80 TgIST-3xHA strain (as constructed in Fig. 1A). Host nuclear TgIST remains significantly elevated above uninfected values at 3, 4, and 5 days p.i. after a 24hr 50nM rapamycin pulse exposure 1day post-infection. Measurements were made from three replicate experiments at each time point from infected nuclei containing a single vacuole at each timepoint. Mean gray values were normalized to uninfected nuclei. Symbol conventions were used as in Figure 2 to indicate significant increases from uninfected values.

Asterisks (*) indicate a statistically significant increase (**** p < 0.0001, *** p < 0.001, ** p < 0.01, * p< 0.05) compared to the uninfected group, whereas hashtag symbols (#) indicate a statistically significant decrease (#### p < 0.0001, ### p < 0.001, ## p < 0.01, # p < 0.05) in infected groups compared to the 1 day post-infection time point.

**(B)** Representative images of a bradyzoite differentiation and nuclear TgIST intensity at 3, 4, and 5 days p.i. Bradyzoite differentiation was assessed by SalmonE antibody to glycosylated CST1 (green), nuclei labeled with DAPI (blue), and TgIST-3xHA labeled with HA antibody. Infected host cell nuclei are indicated by dashed ovals. Scale bar equals 10 µm.

